# Predictive functionality of bacteria in naturally fermented milk products of India using PICRUSt2 and Piphillin pipelines

**DOI:** 10.1101/2021.08.06.455378

**Authors:** H. Nakibapher Jones Shingling, Jyoti Prakash Tamang

**Affiliations:** DAICENTER (DBT-AIST International Centre for Translational and Environmental Research) and Bioinformatics Centre, Department of Microbiology, School of Life Sciences, Sikkim University, Gangtok 737102, Sikkim, India

**Keywords:** Metagenome gene prediction, PICRUSt2, Piphillin, naturally fermented milk products, lactic acid bacteria

## Abstract

Naturally fermented milk (NFM) products are popular food delicacies in Indian states of Sikkim and Arunachal Pradesh. Bacterial communities in these NFM products of India were previously analysed by high-throughput sequence method. However, predictive gene functionality of NFM products of India has not been studied. In this study, raw sequences of NFM products of Sikkim and Arunachal Pradesh were accessed from MG-RAST/NCBI database server. PICRUSt2 and Piphillin tools were applied to study microbial functional gene prediction. MUSiCC-normalized KOs and mapped KEGG pathways from both PICRUSt2 and Piphillin resulted in higher percentage of the former in comparison to the latter. Though, functional features were compared from both the pipelines, however, there were significant differences between the predictions. Therefore, a consolidated presentation of both the algorithms presented an overall outlook into the predictive functional profiles associated with the microbiota of the NFM products of India.

## Introduction

Naturally fermented milk (NFM) products are popular food items in daily diets of ethnic people of Arunachal Pradesh and Sikkim in India, which include *dahi, mohi, gheu*, soft-*chhurpi*, hard-*chhurpi, dudh-chhurpi, chhu, somar, maa, philu, shyow, mar, chhurpi*/*churapi, churkam* and *churtang*/*chhurpupu* (Rai et al. 2016; Tamang et al. 2021). Previously, taxonomic analysis using high-throughput sequencing (HTS) of NFM products of Arunachal Pradesh and Sikkim viz. *chhurpi, churkam mar*/*gheu* and *dahi*, have been studied (Shangpliang et al. 2018). We have recorded the abundance of phylum *Firmicutes* with predominated species of lactic acid bacteria (LAB) viz. *Lactococcus lactis* (19.7%) and *Lactobacillus helveticus* (9.6%) and *Leuconostoc mesenteroides* (4.5%) and acetic acid bacteria (AAB): *Acetobacter lovaniensis* (5.8%), *Acetobacter pasteurianus* (5.7%), *Gluconobacter oxydans* (5.3%), and *Acetobacter* syzygii (4.8%) (Shangpliang et al. 2018). Application of shotgun metagenomics is one of the commonly used methods for understanding the microbial-associated gene functional characteristics (Quince et al. 2017). However, alternately functional profiles of a microbial community can also be inferred indirectly by marker-gene surveys such as 16S rRNA gene (Ortiz-Estrada et al. 2019; Bokulich et al. 2020). Bioinformatics pipelines such as Phylogenetic Investigation of Communities by Reconstruction of Unobserved States version2 (PICRUSt2) (Douglas et al. 2020) and Piphillin (Narayan et al. 2020) among others are some of the well-known tools for microbial predictive functionality studies from various NGS-related metagenomic data (Ortiz-Estrada et al. 2019; Bokulich et al. 2020). These pipelines have also been applied in fermented milk products to infer the functional gene predictions (Zhang et al. 2017; Zhu et al. 2018; Chen et al. 2020; Choi et al. 2020a,b). Microbiota present in NFM products harbour probiotic properties and impart several health-promoting benefits to consumers (Bengoa et al. 2019; Tamang et al. 2020; García-Burgos et al. 2020). Predictive gene functionality in NFM products of India has not been analysed yet. Hence, the present study is aimed to predict the microbial functional contents of 16S rRNA gene sequencing data of NFM products of India, previously analysed by high-throughput sequencing method (Shangpliang et al. 2018), using PICRUSt2 and Piphillin pipelines.

## Material and Methods

### Pre-analysis prior to predictive functionality analysis

Raw sequences of NFM products of Arunachal Pradesh and Sikkim in India analysed by HTS method (Supplementary Table 1) were accessed from MG-RAST/NCBI database server and were used in this study. Raw reads were processed using QIIME2-2020.6 (https://docs.qiime2.org/2020.6/) (Bolyen et al. 2019). After importing into QIIME2 environment, Q-score based filtering and denoising was performed using Divisive Amplicon Denoising Algorithm (DADA2) (Callahan et al. 2016) via qiime dada2 denoise-paired plugin. Quality-filtered sequences were then clustered against SILVA v132 (Quast et al. 2012) databases and followed by taxonomic assignment using q2-vsearch-cluster-features-closed-reference (Rognes et al. 2016).

### Predictive functionality analysis

#### PICRUSt2 analysis (https://github.com/picrust/picrust2/wiki)

Quality-filtered clustered sequences were feed into PICRUSt2 algorithm (Douglas et al. 2020) using via q2-vsearch-cluster-features-closed-reference (Rognes et al. 2016). PICRUSt2 deduced the predictive functionality of the marker genes by using a standard integrated genomes database. Firstly, multiple assignment of the exact sequence variants (ESVs) was performed using HMMER (http://www.hmmer.org/). Placements of ESVs in the reference tree with evolutionary placement-ng (EPA-ng) algorithm (Barbera et al. 2019) and Genesis

Applications for Phylogenetic Placement Analyses (GAPPA) omics (Czech and Stamatakis 2019) were applied. Prediction of gene families was run using a default castor R package (Louca and Doebeli 2018) with the default algorthim run (maximum parsimony) and metagenome prediction was acquired using metagenome_pipeline.py (Ye and Doak 2009).

#### Piphillin analysis (https://piphillin.secondgenome.com/)

Additionally, predictive functionality was also inferred using Piphillin (Narayan et al. 2020), a web-server analysis pipeline. DADA2-clustered representative sequences (.fasta) and abundance frequency table (.csv) were used as inputs for the analysis.

### Statistical analysis and data visualization

Unnormalized Kyoto Encyclopaedia of Genes and Genomes (KEGG) ortholog (KO) profiles of PICRUSt2 and Piphillin predictive were normalized using Metagenomic Universal Single-Copy Correction (MUSiCC) (Manor and Borenstein 2015). The output features were then mapped to KEGG database for systematic analysis of gene functions (Kanehisa et al. 2012). Relative abundance at the category level was plotted as stacked bar-plot using MSEXCEL v365. Statistical analysis for significant features (pathways) was carried out using STAMP (Parks et al. 2014). Normalized predictive features were log-transformed and the differences between PICRUSt2 and Piphillin predictive features were calculated using White’s non-parametric with Benjamini-Hochberg FDR (false discovery rate) (Parks et al. 2014). Non-parametric Spearman’s correlation of the bacteria and functionality was analyzed through Statistical Package for the Social Sciences (SPSS) v20 and the heatmap representation was plotted using ClustVis (Metsalu and Vilo 2015).

## Results

### Microbial predictive gene functionality

A total of 1109 error-corrected ESVs was obtained from DADA2 analysis and about 268 SILVA-clustered sequences were used for the downstream predictive analysis. A total of 5995 MUSiCC-normalized KOs and 181 mapped KEGG pathways was obtained from PICRUSt2 analysis. Similarly, a total of 5245 MUSiCC-normalized KOs and 157 mapped KEGG pathways was obtained from Piphillin analysis. Overall, both PICRUSt2 and Piphillin pipelines showed a similar pattern (Fig. 1), except in the metabolism category where the PICRUSt2 was significantly higher in comparison to that predicted by Piphillin pipeline (Fig. 2). Additionally, at the super pathway level, PICRUSt2 prediction showed significantly high in amino acid metabolism, metabolism of cofactors and vitamins, energy metabolism, and biosynthesis of other secondary metabolites (Fig. 2). On the other hand, predictive super pathways which included carbohydrate metabolism, xenobiotics biodegradation and metabolism, metabolism of other amino acids, lipid metabolism, metabolism of terpenoids and polyketides, glycan biosynthesis and metabolism, and nucleotide metabolism were significantly higher through Piphillin prediction (Fig. 2). Significant metabolic-related pathways inferred by both PICRUSt2 and Piphillin tools were compared showing several functional features predicted by these two pipelines (Fig. 3).

**Figure 1:**
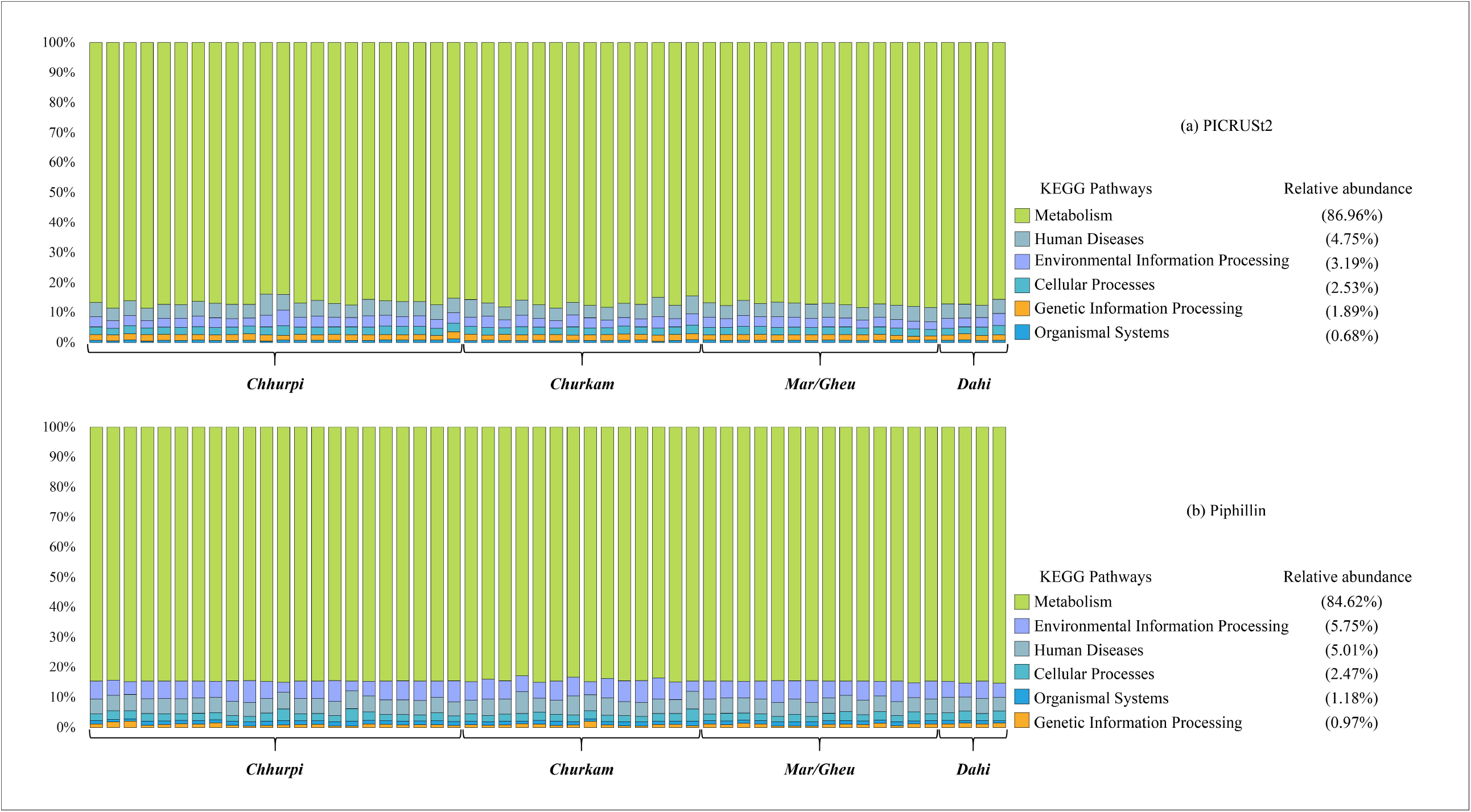
An overall categorical representation of the MUSiCC-normalized predictive microbial functions as inferred by (a) PICRUSt2 and (b) Piphillin.

**Figure 2:**
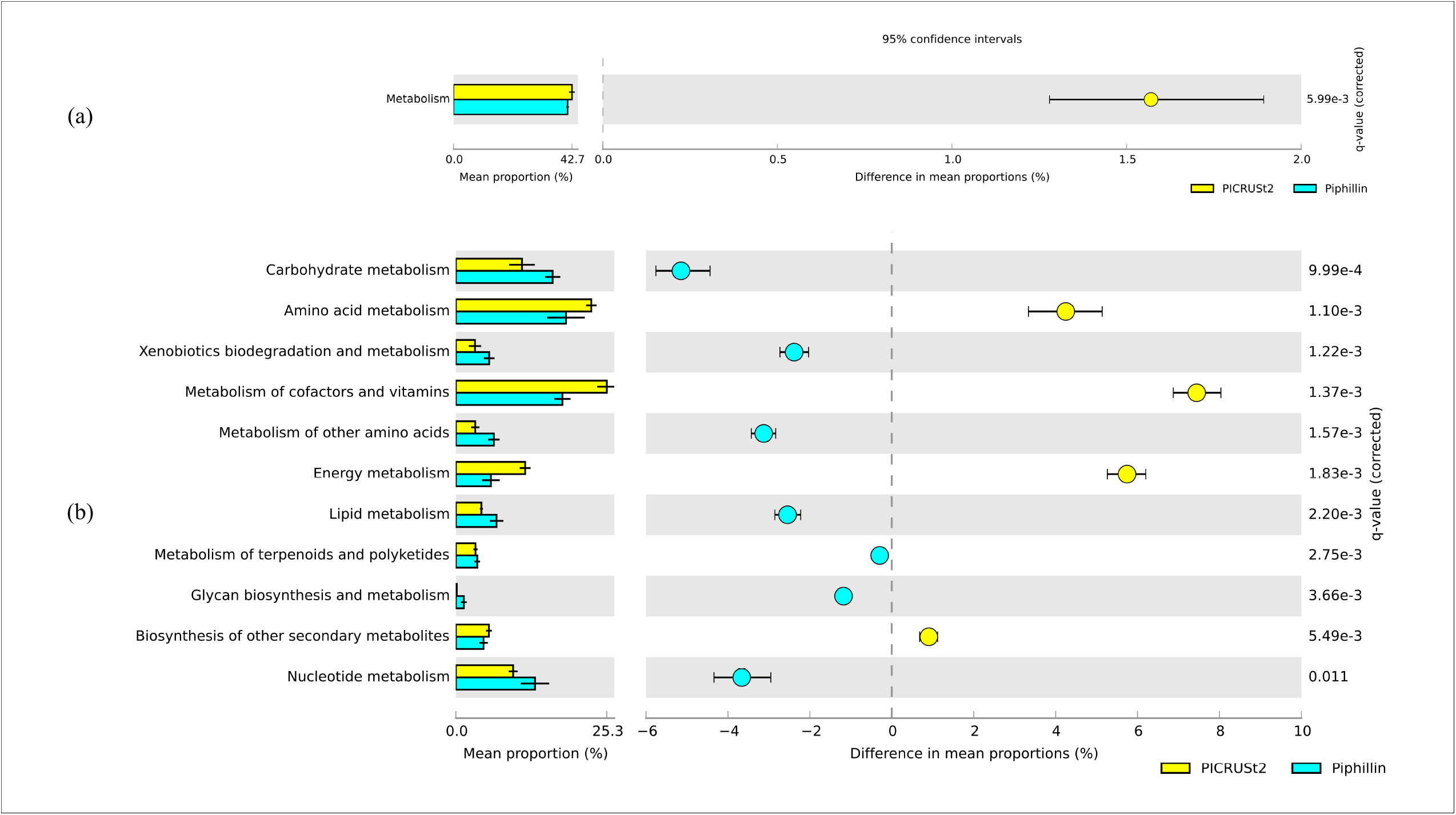
Extended error bar chart representation of the significant predictive functionalities as inferred by both PICRUSt2 and Piphillin. (a) Overall, metabolism is significantly higher in PICRUSt2 analysis as compared to that of Piphillin, however, (b) a shared difference was observed at the super-pathway level. Significance (q-value>0.05) was calculated using White’s non-parametric test with Benjamini-Hochberg FDR (false discovery rate) in STAMP.

**Figure 3:**
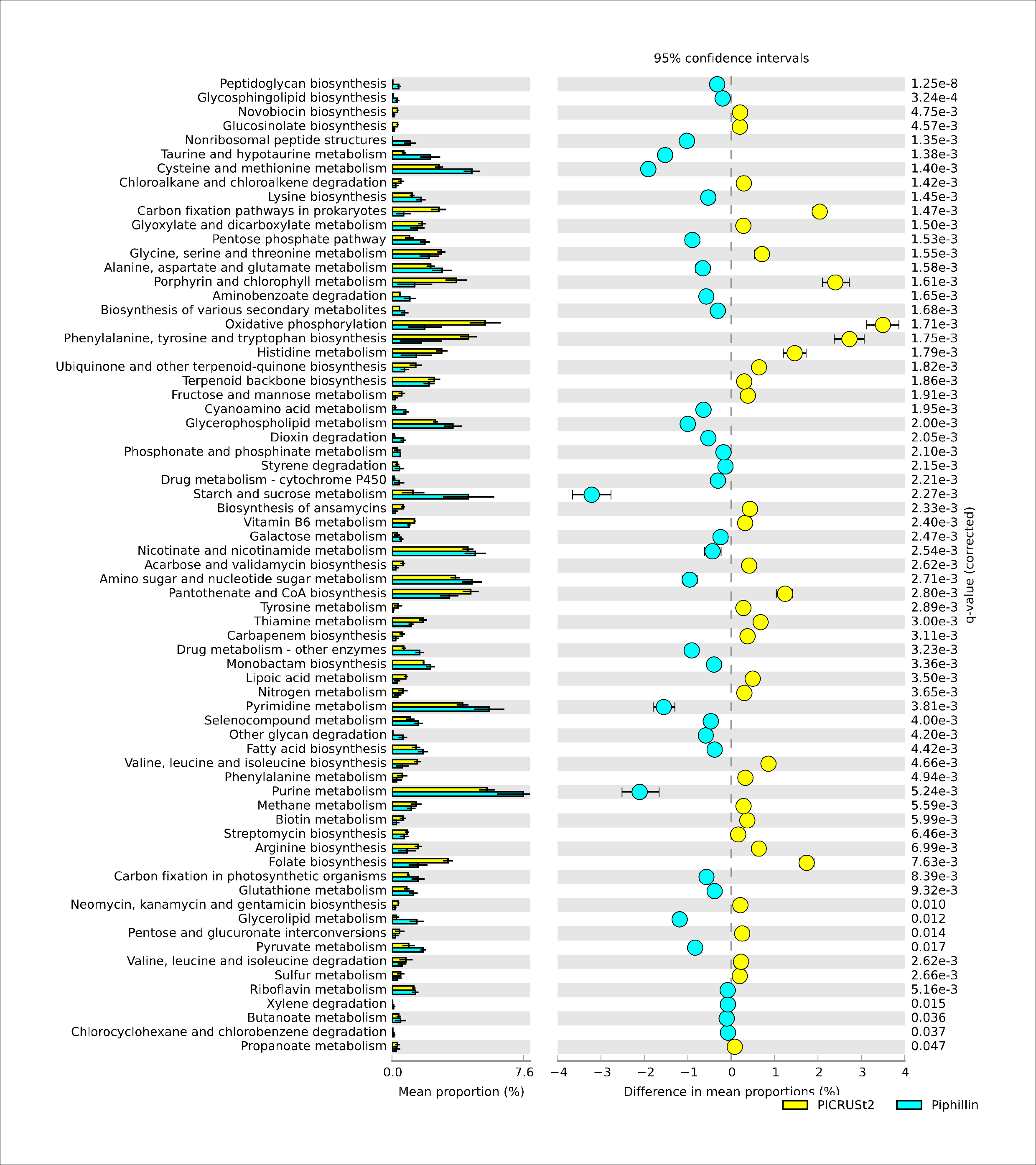
An overall comparison of the significant metabolic pathways as inferred by PICRUSt2 and Piphillin depicting a significant number of functional features predicted by these two pipelines. Significance (q-value>0.05) was calculated using White’s non-parametric test with Benjamini-Hochberg FDR (false discovery rate) in STAMP.

### Non-parametric correlation of bacteria with predictive functionality

Non-parametric Spearman’s correlation analysis resulted in a complex bacterial-functions interaction. *Lactococcus* showed a significant negative correlation with glycerolipid metabolism and ubiquinone and other terpenoid-quinone biosynthesis. *Lactobacillus* showed significant negative correlation with tryptophan metabolism, galactose metabolism, and lipoic acid metabolism while it was observed to be positively significantly correlated with sulphur metabolism. On the other hand, valine, leucine and isoleucine degradation, arginine biosynthesis and ubiquinone and other terpenoid-quinone biosynthesis was positively correlated with *Leuconostoc*, and negatively correlated with galactose metabolism. Furthermore, a significant negative correlation was observed between *Acetobacter* with pathways-tryptophan metabolism, valine, leucine and isoleucine biosynthesis, and lipoic acid metabolism. *Gluconobacter* also showed a significant negative correlation with phenylalanine metabolism, pentose and glucuronate interconversions, fructose and mannose metabolism, and nitrogen metabolism. Glycerolipid metabolism and ubiquinone and other terpenoid-quinone biosynthesis showed significant positive correlation with *Staphylococcus*, which significantly negatively correlated with propanoate metabolism. *Pseudomonas* showed significant negative correlation with fructose and mannose metabolism and significant positive correlation with tyrosine metabolism, valine, leucine and isoleucine degradation, arginine and proline metabolism, galactose metabolism, ubiquinone and other terpenoid-quinone biosynthesis and glutathione metabolism. Additionally, a significant positive correlation was observed between *Acinetobacter* with phenylalanine metabolism, streptomycin biosynthesis, ascorbate and aldarate metabolism, propanoate metabolism, nitrogen metabolism, and biosynthesis of ansamycins (Fig. 4).

**Figure 4:**
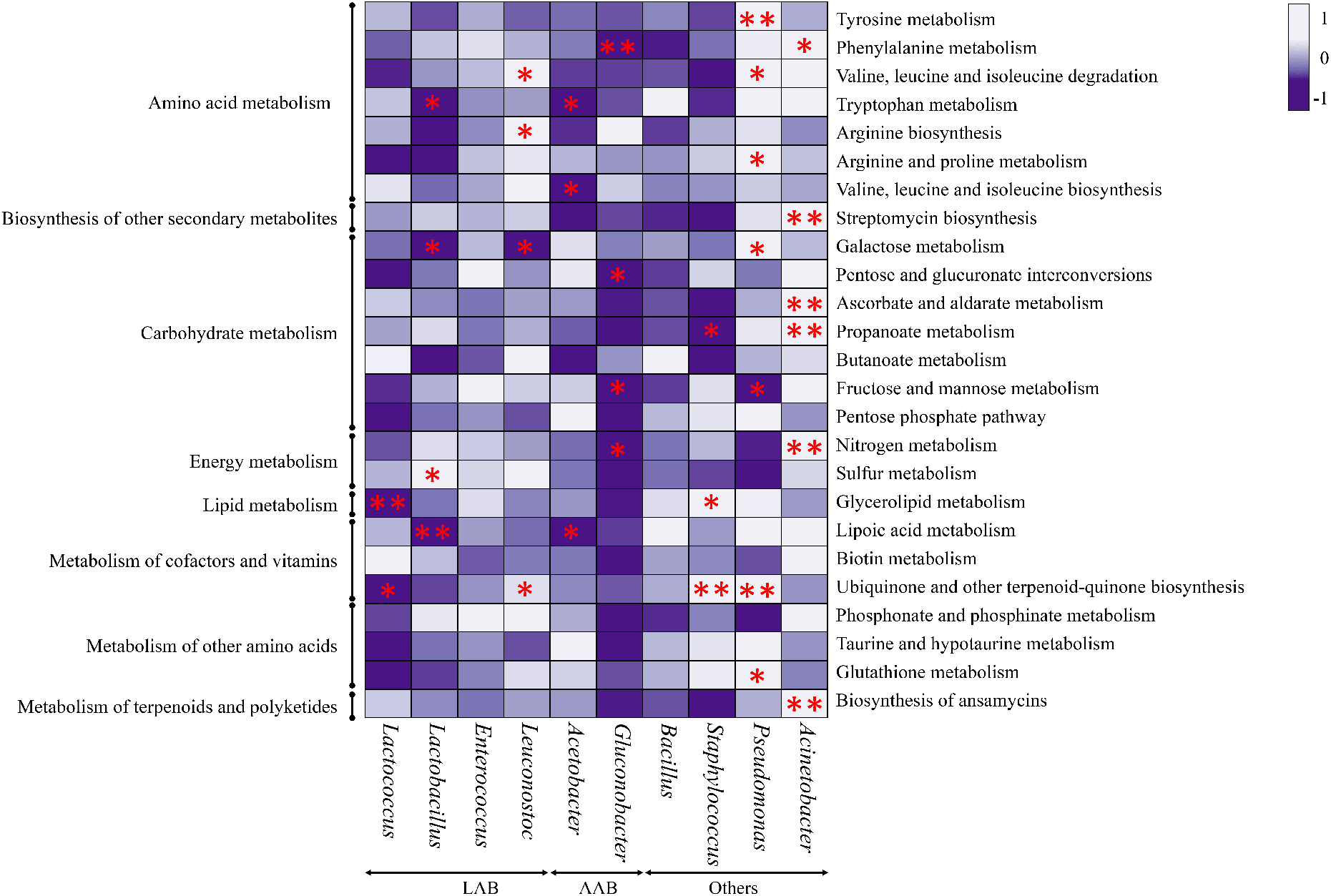
Non-parametric Spearman’s correlation of the ASV-associated predominant bacterial genera of the NFM products with a consolidated functional feature as inferred by both PICRUSt2 and Piphillin. Here, calculation was carried out using Statistical Package for the Social Sciences (SPSS) v20 and heatmap was generated using ClustVis. All significant correlation pairs are denoted by * (*<0.05 and **<0.01). LAB-lactic acid bacteria; AAB-acetic acid bacteria.

## Discussion

In this study, microbial predictive gene functional analysis from targetted-16S rRNA gene was explored using PICRUSt2 and Piphillin pipelines. Inference of predictive functionality using these two said pipelines showed a high metabolism rate, since most of these products are consortia of many metabolically active microbiota (Shangpliang et al. 2018). These findings are similar to recent studies reported from fermented dairy products (Zhang et al. 2017; Zhu et al. 2018; Chen et al. 2020; Choi et al, 2020a,b). The association of various metabolic pathways such as amino acid metabolism, carbohydrate metabolism, energy metabolism, lipid metabolism, metabolism of cofactors and vitamins, and other secondary metabolites with the bacterial genera indicated an active interaction of bacteria-function complexity. LAB are predominant microbiota in many ethnic fermented milk products of India followed by few AAB (Tamang et al. 2000; Dewan and Tamang 2006, 2007; Shangpliang et al. 2018; Ghosh et al. 2019; Shangpliang and Tamang 2021). Spearman’s correlation of the predominant bacterial genera with the predictive functionality resulted in a complex microbial-functions interaction in NFM products of Sikkim and Arunachal Pradesh. Metabolic activity such as amino acid metabolism is important in dairy products as they contribute in development of flavour (Yvon and Rijnen 2001). Similarly, carbohydrate metabolism does also play a major role in flavour and aroma development in milk fermentation (Pan et al. 2014). The abundance of functional pathways related to metabolism of amino acids, lipid, energy and carbohydrates were earlier reported in fermented milk and milk products (Zhang et al. 2017; Ramezani et al. 2017; Zhu et al. 2018; Yasir et al. 2020; Chen et al. 2020). A high correlation of functional properties and LAB have also been reported in cheeses (Yang et al. 2020), since LAB are the most predominant microorganisms in fermented milk products (Rezac et al. 2018; Chen et al. 2020). We observed a positive correlation of *Staphylococcus* with the predictive metabolic features of these NFM products, and interestingly, *Staphylococcus* is metabolically active in dairy products playing functional activities such as amino acid metabolism, carbohydrate metabolism, lipid metabolism and nitrogen metabolism (Leroy et al. 2020). We also observed the presence of significant correlation of bacteria with cofactors and vitamins metabolism such as ubiquinone and other terpenoid-quinone biosynthesis and lipoic acid metabolism, which are essential for other microbial metabolism (Yao et al. 2020). Apart from LAB, AAB have also contributed to many functional features in NFM products; AAB involve in protein metabolism, production of secondary metabolites and volatile compounds (Illeghems et al. 2015; Ai et al. 2019).

Functional profiles from both PICRUSt2 and Piphillin were normalized using MUSiCC (Manor and Borenstein 2015), which is a marker gene-based method which use universal single-copy genes for biasness correction of gene abundances (Noecker et al. 2017). Normalization using MUSiCC have proven necessary for gene functional studies (Vincent et al. 2017), rescaling the abundant predicted KOs to the actual average gene copy number, correcting several known biases (Manor and Borenstein 2017). Piphillin is usually applied in human clinical samples (Iwai et al. 2016); whereas PICRUSt2 is widely used for environmental samples (Douglas et al. 2020). However, these pipelines have also been widely used in dairy products (Choi et al. 2020a,b).

From our present analysis, PICRUSt2 analysis generated more predicted KOs and KEGG pathways in comparison to that of Piphillin. Though, significant differences were observed, however, there are functions which were predicted only from PICRUSt2 and missing in Piphillin and vice versa. Therefore, consolidated predictive functions from both these pipelines are necessary for a comprehensive outlook into the potential of bacteria associated with NFM products. Though predictive functionality study of the microbiota associated with NFM products at present is only speculations using bioinformatics tools, a general outlook into the potentiality of functions may be studied and compared. Nonetheless, in the absence of shotgun metagenomics data, using PICRUSt2 and Piphillin serves to be the reliable analysis for microbial predictive gene function.

## Conclusion

Bacterial community in NFM products showed many functional features with many important health benefits to consumers. We applied PICRUSt2 and Piphillin tools to infer the predictive functional features of microbiota associated with the ethnic fermented milk products of India. Therefore, such studies may be used for future comparison with detailed gene functionality studies of other fermented foods elsewhere.

## Acknowledgements

The authors are grateful to the Department of Biotechnology, Govt. of India through the DAICENTER project. HNJS is grateful to DBT for Junior Research Fellowship.

## Funding

This current research is supported by Department of Biotechnology, Govt. of India through DBT-AIST International Centre for Translational and Environmental Research (DAICENTER) project.

## Authors’ contributions

HNJS did analysis and bioinformatics analysis. JPT has supervised the bioinformatics analysis and finalised the manuscript.

## Availability of data and materials

Raw sequences were accessed from MG-RAST server having the MG-RAST ID number 4732361 to 4732414. The same were accessed from NCBI database server under the BioProject No. PRJNA661385 with accession numbers SAMN16056817 to SAMN16056870.

## Declaration of Competing Interest

The authors declare that they have no competing interests.

## Notes

### Competing Interest Statement

The authors have declared no competing interest.

