## Supplementary table 1 for "Predictive functionality of bacteria in naturally fermented milk products of India using PICRUSt2 and Piphillin pipelines"

**Supplementary Table 1:** Bacterial genera identified from different NFM products of Sikkim and Arunachal Pradesh (Shangpliang et al. 2018).

| **Genus** | **Region** | **Relative percentage of abundance (%)** | | | | | | | | | | |
| --- | --- | --- | --- | --- | --- | --- | --- | --- | --- | --- | --- | --- |
|  |  | ***Lactococcus*** | ***Lactobacillus*** | ***Acetobacter*** | ***Leuconostoc*** | ***Pseudomonas*** | ***Staphylococcus*** | ***Gluconobacter*** | ***Bacillus*** | ***Acinetobacter*** | ***Enterococcus*** | **Others (<1%)** |
| *Chhurpi* | Sikkim | 30.54 | 19.18 | 15.9 | 12.7 | 6.04 | 4.06 | 2.45 | 1.66 | 0.71 | 1.28 | 5.48 |
| *Gheu* | Sikkim | 28.92 | 4.43 | 4.51 | 4.00 | 49.81 | 0.45 | 1.13 | 0.24 | 2.15 | 0.21 | 4.15 |
| *Dahi* | Sikkim | 42.29 | 13.49 | 9.32 | 17.68 | 9.57 | 0.86 | 2.56 | 0.55 | 1.59 | 0.19 | 1.90 |
| *Chhurpi* | Arunachal Pradesh | 51.06 | 15.80 | 10.87 | 5.10 | 3.88 | 4.43 | 2.27 | 2.26 | 0.77 | 0.90 | 2.66 |
| *Churkam* | Arunachal Pradesh | 38.98 | 14.81 | 11.95 | 5.05 | 3.46 | 13.16 | 2.36 | 3.16 | 1.29 | 1.38 | 4.40 |
| *Mar* | Arunachal Pradesh | 35.22 | 14.12 | 19.22 | 8.65 | 4.56 | 4.65 | 7.11 | 2.56 | 0.82 | 0.98 | 2.11 |
|  | Average (%) | 37.84 | 13.64 | 11.96 | 8.86 | 12.89 | 4.60 | 2.98 | 1.74 | 1.22 | 0.82 | 3.45 |
